# A data standard for the reuse and reproducibility of any stable isotope probing-derived nucleic acid sequence (MISIP)

**DOI:** 10.1101/2023.07.13.548835

**Authors:** Abigayle Simpson, Elisha M. Wood Charlson, Montana Smith, Kathleen Beilsmith, Ben Koch, Ramona L. Walls, Roland C. Wilhelm

## Abstract

DNA/RNA-stable isotope probing (SIP) is a powerful tool to link *in situ* microbial activity to sequencing data. Every SIP dataset captures distinct information about microbial community metabolism, kinetics, and population dynamics, offering novel insights according to diverse research questions. Data re-use maximizes the information available from the time and resource intensive SIP experimental approach. Yet, a review of publicly available SIP sequencing metadata reveals that critical information necessary for reproducibility and reuse is often missing. Here, we outline the Minimum Information for any Stable Isotope Probing Sequence (MISIP) according to the Minimum Information for any (x) Sequence (MIxS) data standard framework and include examples of MISIP reporting for common SIP approaches. Our objectives are to expand the capacity of MIxS to accommodate SIP-specific metadata and guide SIP users in metadata collection when planning and reporting an experiment. The MISIP standard requires five metadata fields: isotope, isotopolog, isotopolog label and approach, and gradient position, and recommends several fields that represent best practices in acquiring and reporting SIP sequencing data (*ex.* gradient density and nucleic acid amount). The standard is intended to be used in concert with other MIxS checklists to comprehensively describe the origin of sequence data, such as for marker genes (MISIP-MIMARKS) or metagenomes (MISIP-MIMS), in combination with metadata required by an environmental extension (*e.g.*, soil). The adoption of the proposed data standard will assure the reproducibility and reuse of any sequence derived from a SIP experiment and, by extension, deepen understanding of *in situ* biogeochemical processes and microbial ecology.

## Introduction

The invention of DNA/RNA-stable isotope probing (SIP) was a groundbreaking achievement that continues to advance our knowledge of microbiology, microbial ecology, and biogeochemistry [1, 2]. SIP provides a method to link sequencing data with microbial activity based on the incorporation of an isotopically labeled compound of interested (*isotopolog)* into the nucleic acids of active populations (Figure 1A). Subsequent innovations have improved the utility of SIP for quantifying the differential activity of populations within whole microbial communities [3–5]. To achieve this, a typical SIP experiment generates large amounts of sequencing data due to the necessity of sampling multiple density gradient fractions and the use of paired controls (Figure 1B). Although there are fundamental similarities in the metadata generated during SIP experiments, most publicly archived SIP sequence data lack sufficient metadata (Figure 2A), hindering the reproducibility of results and data reuse.

**Figure 1.**
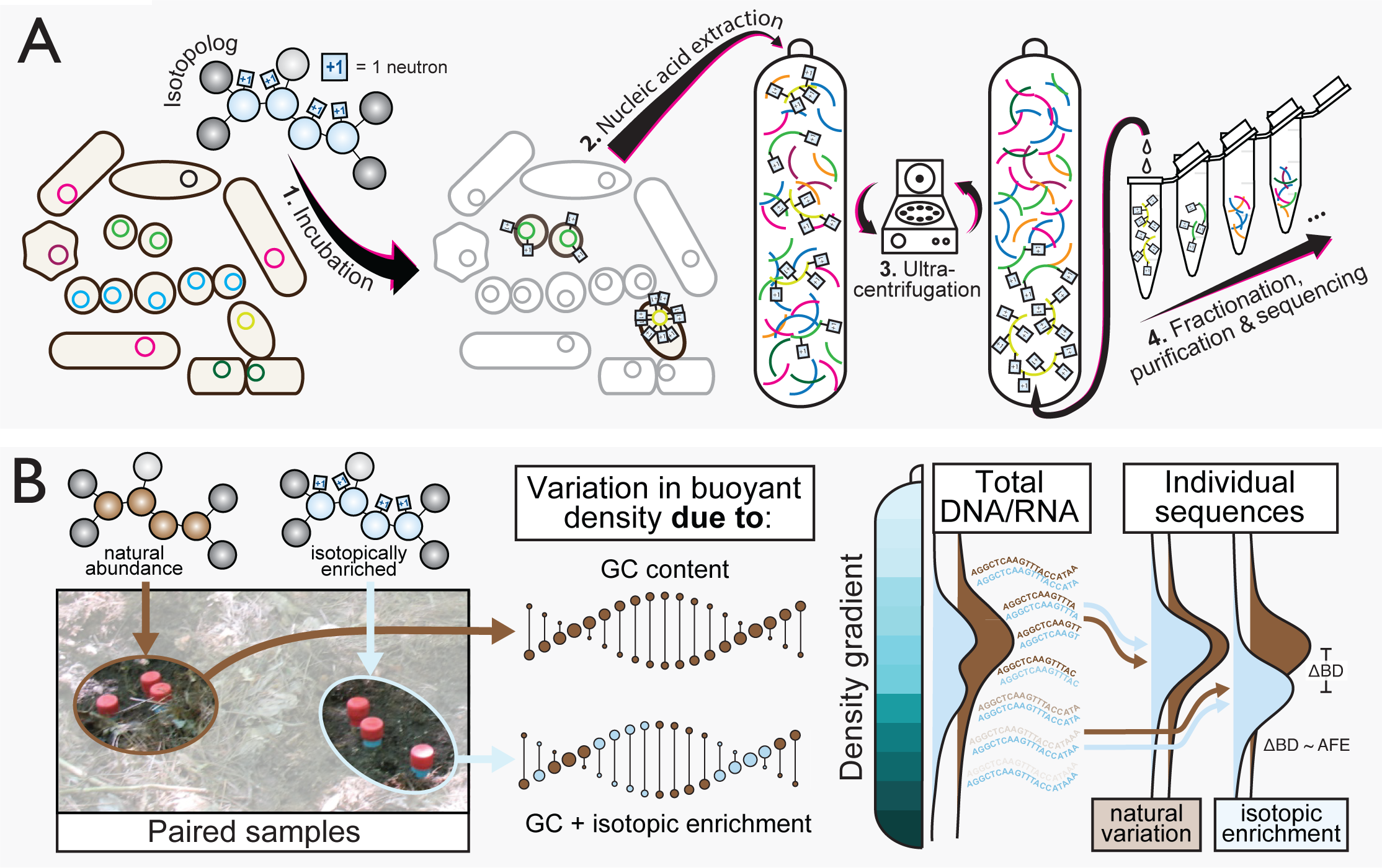
An overview of the key principles of the DNA/RNA-SIP method, illustrating the impact of density gradient separation on sequencing data composition (**Panel A**) and the need for paired samples supplied with different isotopologs to distinguish natural variation in buoyant density from the effects of isotopic enrichment (**Panel B**). Panel A shows the standard series of steps performed to isotopically enrich and separate nucleic acids using density gradient (‘isopycnic’) ultracentrifugation. In this case, an isotopically enriched isotopolog is incubated in the presence of a microbial community, during which the nucleic acids of active cells incorporate artificially high concentrations of the isotope (step 1). Whole nucleic acids are extracted from the sample (step 2) and centrifuged at high force to establish a density gradient and separation of nucleic acids based on buoyant density (step 3). The final step is to fractionate the separated nucleic acids, followed by purification and sequencing (step 4). Panel B depicts the use of paired samples supplied with either natural abundance (‘unlabeled’) or isotopically enriched isotopologs to isolate the variation in buoyant density of DNA/RNA, due to GC content, from the effects of isotopic enrichment. The atom fraction excess of isotope (AFE) can be estimated on a per sequenced basis using the change in buoyant density (ΔBD) between paired samples [3], demonstrating the importance of paired samples.

**Figure 2.**
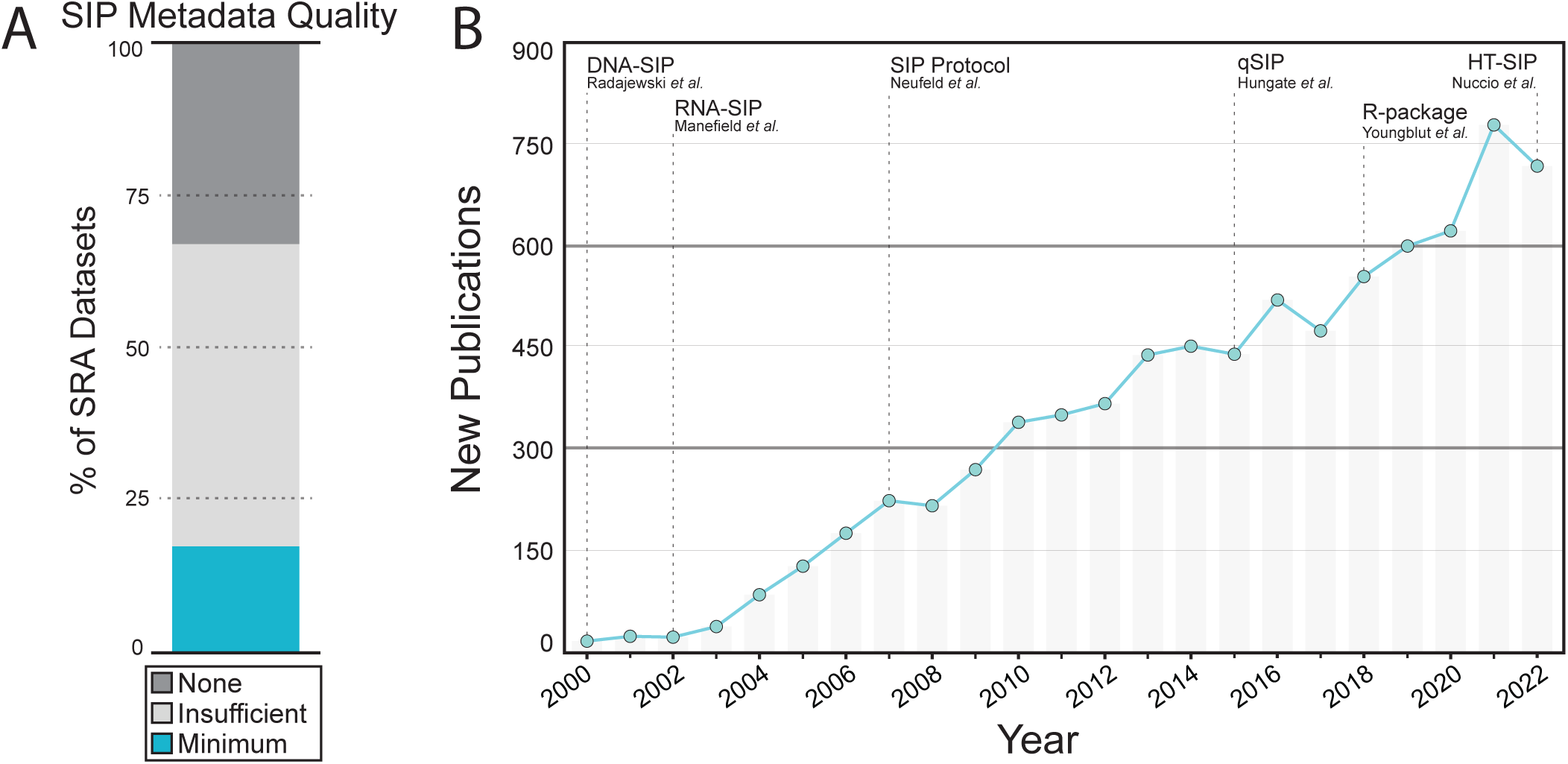
A summary of the quality of metadata associated with SIP sequencing data available in the Sequence Read Archive (A) and the growth in the number of DNA/RNA-SIP studies published over time (B). In (A), an overview of the metadata quality of studies with SRAs containing more than five samples. SRAs which reported the isotopolog, isotopolog label status, and gradient position for each entry met the “Minimum” requirements, while those that reported at least one of these items were “Insufficient”, and those that reported none were categorized as “None.” In (B), the new number of published studies by year was identified using the search term: “Stable Isotope Probing” and “DNA” or “RNA” in Google Scholar. A timeline showing the major advances in the SIP methodology was included.

The need to facilitate metadata standardization of SIP sequencing data is growing, as the number of studies generating SIP sequencing data has been growing year-upon-year (Figure 2B), with further growth expected due to improvements in automated sample processing [6]. Furthermore, the reuse of SIP sequence data has considerable value given the expense and labor involved in these experiments and the information gained by extrapolating across various isotopologs or studies [7–9]. Here, we propose a minimum set of required metadata fields for SIP-derived sequencing data, as well as a recommended set that embodies the best practices in acquiring and reporting SIP sequencing data.

### Essential properties of SIP sequence data

In all cases, sequence data generated from a SIP experiment originates from nucleic acid pools that have been fractionated by isopycnic (or ‘density gradient’) separation [10]. The gradient separates nucleic acids based on differences in buoyant density due to the added mass per nucleic acid from the incorporation of heavy stable isotopes (*ex.* ^13^C, ^15^N, ^2^H, or ^18^O) during the metabolism of an isotopically labeled source isotopolog (*ex.* ^13^CO_2_, ^15^NH ^+^, or H ^18^O). The fractionated pools of nucleic acids are then sequenced to resolve differences in buoyant densities corresponding to isotopic enrichment. Many fractions may be sequenced to determine fine-scale isotopic enrichment (‘density-resolved SIP’), or fractions may be pooled before sequencing to compare bulk differences in buoyant density. Either way, each nucleic acid sample typically generates multiple sequencing libraries.

There is natural variation in the buoyant density of nucleic acids due to the effect of GC content on genome density [11]. This variability creates sample-specific buoyant density distributions of nucleic acids based on the genomic composition of the biological community under study. To control for this, a standard SIP approach involves comparing sequence data generated from identically treated sample pairs: one that received an unlabeled isotopolog (*i.e.*, natural abundance) and another that received an artificially labeled isotopolog [10]. The reproducibility and reuse of SIP sequence data cannot be achieved without information about the position from which the nucleic acids originated in the density gradient and the sample pairing, as well as information about the stable isotope(s) and source isotopolog compound(s) used.

### The case for a SIP-specific data standard

At present, there is no convention for the handling of SIP sequence metadata despite the archival of hundreds of datasets in public databases, spanning over two decades of sequencing types (clone libraries to shotgun metagenomes) from diverse environments (Figure S1). A formal standard describing the minimum information for any SIP sequence is needed to ensure the adoption of FAIR principles (findability, accessibility, interoperability, and reusability) for data reuse [12]. Currently, there is no stable identifier for SIP sequence data, requiring users to glean this information from the study description or associated publication, which poses challenges to findability. Additionally, the absence of data labeling requirements creates the risk of naïve users misinterpreting ambiguously labeled SIP data and failing to account for biases caused by gradient fractionation. Furthermore, there is no common vocabulary for the diverse types of SIP sequence metadata generated during the course of an experiment. The formalization of a specific, richly described, and consistent vocabulary is critical for interoperability among SIP sequence data. Furthermore, prioritizing the inclusion of several metadata fields will greatly improve the machine readability, analyses, and interpretation of SIP sequence data.

For these reasons, we propose a SIP data standard (MISIP) designed to be combined with existing ‘minimum information about a marker gene sequence’ [13] (‘MISIP-MIMARKS’) and the ‘minimum information about a metagenome sequence’ [14] (‘MISIP-MIMS’) checklists developed by the Genomic Standards Consortium (GSC) [15]. MISIP will expand the capacity to accommodate SIP-specific metadata in sequence archives and guide the archiving of critical metadata. The standard includes a required and a recommended set of metadata fields, though inclusion of the recommended set is not mandated (Table 1). In the following sections, we provide descriptions of the metadata fields and justify their designation as required or recommended in the associated sequencing archival checklist. We intend this information to serve as a reference for users of MISIP and to help newcomers to SIP experiments collect and curate better metadata.

**Table 1.**
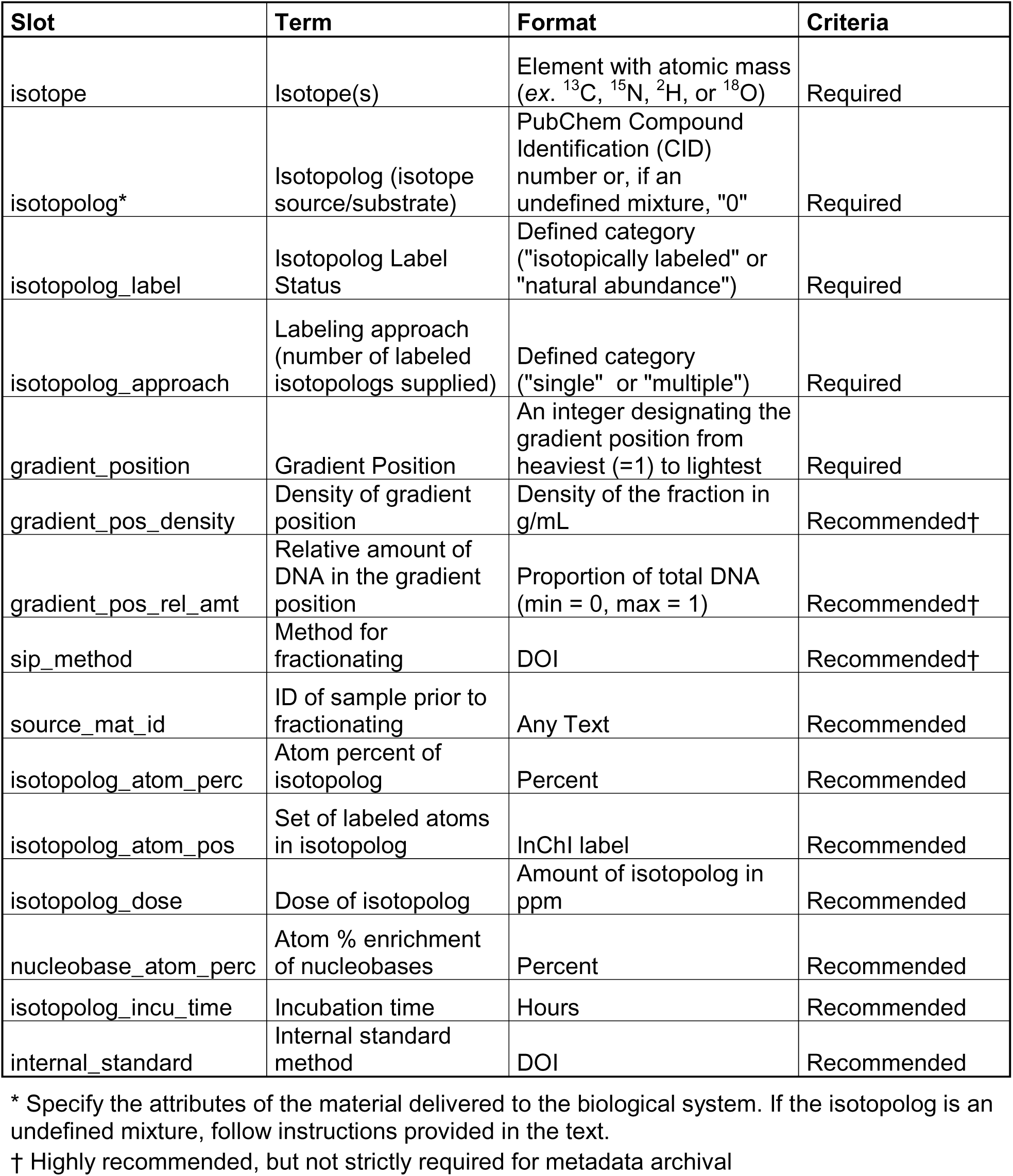
The minimum set of SIP-specific metadata required or recommended by the MISIP data standard. The table summarizes the standardized vocabulary, data format, and whether it is required or recommended (*i.e.*, optional). Descriptions of each metadata field and justifications for their inclusion are provided in the text.

| Slot | Term | Format | Criteria |
| --- | --- | --- | --- |
| isotope | Isotope(s) | Element with atomic mass (ex. <sup>13</sup>C, <sup>15</sup>N, <sup>2</sup>H, or <sup>18</sup>O) | Required |
| isotopolog* | Isotopolog (isotope source/substrate) | PubChem Compound Identification (CID) number or, if an undefined mixture, "0" | Required |
| isotopolog_label | Isotopolog Label Status | Defined category ("isotopically labeled" or "natural abundance") | Required |
| isotopolog_approach | Labeling approach (number of labeled isotopologs supplied) | Defined category ("single" or "multiple") | Required |
| gradient_position | Gradient Position | An integer designating the gradient position from heaviest (=1) to lightest | Required |
| gradient_pos_density | Density of gradient position | Density of the fraction in g/mL | Recommended† |
| gradient_pos_rel_amt | Relative amount of DNA in the gradient position | Proportion of total DNA (min = 0, max = 1) | Recommended† |
| sip_method | Method for fractionating | DOI | Recommended† |
| source_mat_id | ID of sample prior to fractionating | Any Text | Recommended |
| isotopolog_atom_perc | Atom percent of isotopolog | Percent | Recommended |
| isotopolog_atom_pos | Set of labeled atoms in isotopolog | InChI label | Recommended |
| isotopolog_dose | Dose of isotopolog | Amount of isotopolog in ppm | Recommended |
| nucleobase_atom_perc | Atom % enrichment of nucleobases | Percent | Recommended |
| isotopolog_incu_time | Incubation time | Hours | Recommended |
| internal_standard | Internal standard method | DOI | Recommended |

\* Specify the attributes of the material delivered to the biological system. If the isotopolog is an undefined mixture, follow instructions provided in the text.
† Highly recommended, but not strictly required for metadata archival

### The minimum information for any SIP-derived sequence

The MISIP data standard requires metadata fields that are essential for the reuse and reproducible analysis of SIP data. These fields are not currently accommodated by any MIxS checklists. MIxS checklists are sets of required, recommended, and optional metadata terms used to describe different types of sequencing methods that are used by all major sequencing repositories around the world. The SIP-specific fields outlined must be combined with a checklist that describes library preparation and sequencing, such as amplicon (MIMARKS) or shotgun (MIMS) sequence data, plus a MIxS environmental extension that describes the environmental source from which nucleic acids were extracted (*e.g.*, water, soil, or host-associated). All proposed field names are unique and non-redundant according to queries of all current checklists and packages described by the GSC. We have provided examples of SIP-specific metadata curated according to the MISIP standard, including examples of a common gradient pooling approach [16] (Table S2), a more complex SIP experiment [8] (Table S3), and a qSIP approach (Table S4), along with several others examples (Tables S5-S7).

MISIP was developed after an extensive review of existing SIP literature and in conversations with the SIP research community. The standard then underwent iterative refinement with the Compliance and Interoperability Working group of the GSC. The GSC provides the governance and technical infrastructure for standards related to nucleotide sequencing, biological sampling, and related processes but relies on scientific communities like SIP researchers to request and define terms and new checklists or environmental extensions. To increase usability, reproducibility, sustainability, and consistency, GSC switched to managing MIxS standards via an open GitHub repository [17] and using LinkML tooling [18] with the release of MIxS v.6 last year. The MISIP checklists will be included as part of the upcoming MixS v.7 release (Fall 2023), at which point they will be integrated with International Nucleotide Sequence Database Collaboration (INSDC) and other databases. As a living standard, MIxS changes regularly, with stable releases about once per year. Once MISIP is released through MIxS, the latest stable authoritative version of the standard will always be available at: https://w3id.org/mixs.

### Descriptions of required MISIP fields

#### Isotope

The specific stable isotope(s) supplied to the biological system is required information, since the stoichiometry of each element can influence the magnitude of shift in buoyant density of the nucleic acids used to generate SIP sequencing data. For example, fully ^13^C-labeled DNA would produce a larger increase in buoyant density compared to fully ^15^N-labeled DNA, owing to the unequal ratio of approximately 5 carbons to 1 nitrogen in DNA. The ***isotope*** field specifies the element and mass number (*ex*. ^18^O recorded as ‘18O’) of the stable isotope of interest. This field will correspond to the same stable isotope regardless of whether the concentration of the isotope was artificially enriched (*ex.* 99 atom % ^18^O) or occurred at natural abundance (∼0.205 atom % ^18^O), often referred to as a ‘control.’

#### Isotopolog

The central aim of a SIP experiment is to link the isotopic enrichment of nucleic acids to the metabolism of an isotopolog source (or ‘substrate’). The chemical properties of the isotopolog determine how to interpret the underlying metabolic activity that produced the isotopic enrichment of nucleic acids, and the associated changes in the composition of sequence data. For example, certain isotopologs, such as H ^18^O, are used to characterize the whole metabolic activity of a biological system [19], while others, such as ^13^C-labeled phenolic acid, are used to target specific metabolic activity [20]. Surprisingly, the number of accessioned SIP experiments that report the isotopolog is low (<30%; Table S1), making it an often overlooked, but essential, attribute of SIP sequencing data. The ***isotopolog*** field specifies the PubChem Compound Identification (CID) number for the isotopolog serving as the isotope source (*ex.* 6255 for maltose). If the isotopolog is an undefined chemical mixture, the *isotopolog* field should list “0” (which is not associated with any CID), and an additional field should be included to describe the isotopolog mixture in text form. In cases where a sample was not amended with any isotopolog (*ex.* an unamended control), the *isotopolog* field should list “none”, and other isotopolog-describing fields should specify “NA.”

#### Isotopolog label

The standard SIP method requires the pairing of samples that have been supplied with either an isotopically labeled or natural abundance (or ‘unlabeled’) isotopolog to account for shifts in buoyant density from variation in the GC content of genomes [10, 11]. The isotopolog label status is essential to determine whether the buoyant density distribution of nucleic acids reflects isotopic labeling or natural variation. The inclusion of paired samples is not required by MISIP, since the gradient distribution of natural abundance nucleic acid fragments can be modeled [21]. However, the ***isotopolog_label*** field is required as it specifies whether the corresponding isotopolog contains the natural abundance (“natural abundance”) or artificially enriched (“isotopically labeled”) concentration of stable isotope.

In cases where one control sample has been amended with several natural abundance isotopologs and is paired with numerous samples supplied with individual isotopically enriched isotopologs (*ex.* [8]), the control sample (value = “natural abundance”) should be replicated multiple times in the metadata with each corresponding isotopolog field changed to match the paired isotopically-labeled isotopolog (see Table S3 for example).

#### Isotopolog labeling approach

Several isotopes and/or isotopologs may serve as the source of the isotopic enrichment of nucleic acids in a SIP experiment. For example, researchers can use a dual-labeling approach, as previously performed with H_2_ ^18^O and ^13^C-glucose [22]. In these cases, multiple isotopes can contribute to shifts in nucleic acid buoyant density, complicating the analysis and interpretation of SIP sequence data. These datasets should be clearly labeled and easily filtered to ensure appropriate data reuse. The ***isotopolog_approach*** field specifies whether the associated SIP experiment utilized a single isotope and isotopolog (*isotopolog_approach* = ‘single’) or multiple isotopes or isotopologs (*isotopolog_approach* = ‘multiple’) within the same sample.

While multiple-labeling approaches make up less than 10% of accessioned SIP studies (Table S1), these experiments complicate the reporting of a number of other fields, such as the isotope, isotopolog, and isotopolog-describing fields. In these cases, users should list values for these fields in a consistent order, separated with a pipe (*e.g.*, *isotope* = ^13^C | ^18^O, *isotopolog* = 5793 | 962).

#### Gradient position

SIP sequencing data is heavily influenced by differences in the buoyant density of nucleic acids recovered at positions across the density gradient. Thus, MISIP requires information about the gradient position from which the sequenced nucleic acids were recovered. The ***gradient_position*** field is specified as a number starting from the densest (‘heaviest’) gradient fraction (=1) moving in sequential order to the least dense fraction (‘lightest’), keeping with the order in which a gradient is typically fractionated. MISIP users must take special care to ensure that the gradient position numbers match between paired isotopically labeled and natural abundance samples (Figure 3A). The inclusion of unfractionated samples, from which fractionated samples were derived, can be denoted with *gradient_position* = –1.

**Figure 3.**
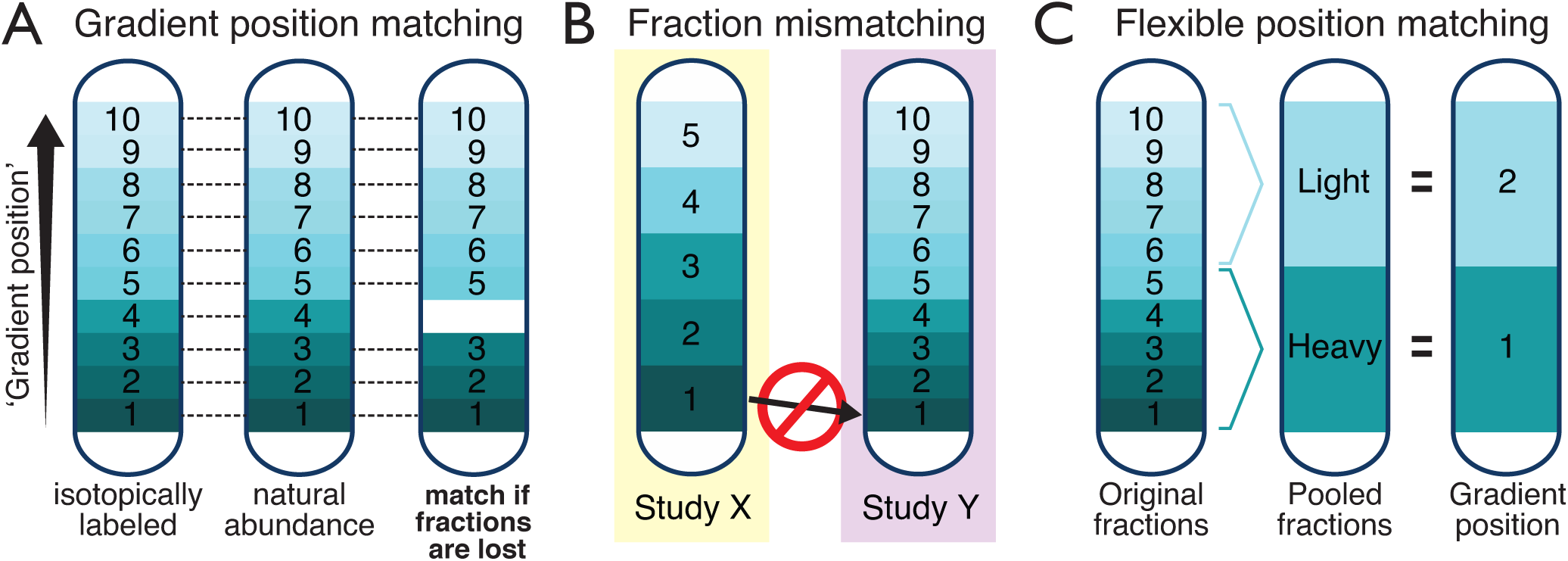
The separation of nucleic acids according to differences in buoyant density due to isotopic enrichment is a defining feature of SIP sequence data. The gradient position designates the location along the density gradient from which the sequenced nucleic acids were recovered. The gradient position generally follows the order in which gradient fractions were collected, accounting for cases where fractions have been lost during processing. Panel A demonstrates the importance of ensuring gradient positions match between paired isotopically labeled and natural abundance samples. Panel B illustrates the disparity in gradient density due to study-specific methods when comparing fraction numbers. Panel C illustrates the use of the gradient position system to accommodate different strategies employed for gradient fraction pooling.

The MISIP standard uses the numerical order of the gradient position to match paired samples because it is the most flexible way to accept SIP sequence data. SIP experiments typically assign either a fraction number or a direct measure of buoyant density to nucleic acids derived from the density gradient. However, the treatment of fractionated nucleic acid pools will depend on the subjective aims of a SIP experiment, leading to diverse and ad hoc ways in which fractions are pooled prior to sequencing. Defaulting to the numerical order for position in the density gradient parallels the common approach of designating fraction number. However, MISIP uses the term ‘gradient position’ to avoid the assumption of equivalence between fractions across studies, since fractionation will yield different volumes and densities depending on methodologies (Figure 3B).

Direct measurement of the buoyant density of each gradient fraction is <u>strongly recommended</u> but not required for several reasons. The measurement of buoyant density, using the refractometric index or by weighing, is not always obtained, nor is it always accurate due to inadequate refractometer calibration or challenges with weighing small volumes (at high precision) caused by evaporation. Furthermore, many studies do not provide information on the methods for converting between refractive index and gradient density, including whether they correct for temperature.

The gradient position is agnostic to the assortment of ways nucleic acids are treated during fractionation. For example, ‘heavy’ and ‘light’ are common designations of the location in the density gradient from which SIP sequence data originates after pooling multiple gradient fractions. In the MISIP gradient position system, these categorical values would be assigned a gradient position of 1 and 2, respectively, which are decoupled from the original fractions numbers (Figure 3C). The gradient position system has its own pitfalls, with positioning becoming discordant when a fraction is lost or when fractionation is inconsistently initiated, though we anticipate the consistency of fractionation will improve over time due to efforts to automate the process [6].

Depositors must take special care to ensure that, at the very least, the gradient position numbers reflect the gradient distribution between paired isotopically labeled and unlabeled controls, should pairs exist in the dataset. Users of MISIP sequence data must also be aware of the potential incongruity between buoyant density and gradient position. These sources of error can be easily resolved by collecting the recommended MISIP metadata, including measurements of gradient density or the use of internal standards to evaluate gradient formation [5].

### Recommended MISIP sequence metadata

The set of required SIP-specific fields in the MISIP standard were chosen to accommodate the fullest diversity of SIP experimental approaches and sequence data, while maintaining the minimum basis for reuse. In most cases, the minimum requirements fall well short of the most useful metadata associated with the nucleic acid pools sequenced during a SIP experiment. As part of the standard, we propose a set of recommended fields and highlight those we consider to be the gold standard (17) for supporting the most sophisticated and quantitative reuse of SIP sequencing data. Example metadata, curated as described below, is available in Tables S2-S7.

#### Gradient position density

⍰A density measurement for each gradient position is essential to (i) evaluate the formation of the density gradient, (ii) normalize among gradient fractionated samples, and (iii) calculate the change in buoyant density (‘ΔBD’) to estimate the degree of isotopic enrichment of sequenced nucleic acids [3]. Measurements of gradient density are easily obtained using a refractometer or analytical balance [10]. Notably, density measurements can be subject to methodological biases due to the small volumes often used in measurement and the inaccuracy of refractometers. The ***gradient_pos*_*density*** field is specified as a numerical value corresponding to the density of gradient solution in grams per milliliter (g · mL^-1^).

#### Relative amount of nucleic acid at gradient position

⍰Measurement of the relative quantity of sequenced nucleic acids in each gradient fraction enables the estimation of a taxon-, genome- or gene-specific isotopic-enrichment, or atom fraction excess (AFE). The method to calculate AFE is termed ‘quantitative SIP (qSIP)’ and, depending on the sequence data type, may require qPCR-based estimates of gene abundance [3, 5]. The ***gradient_pos_rel_amt*** field is specified as the proportion of sequenced nucleic acids relative to the total amount of nucleic acids added to the density gradient prior to ultracentrifugation and must not exceed 1. For shotgun sequencing data, this proportion should be calculated from total nucleic acids (Figure 4A), while for amplicon sequence data, this proportion should be calculated from the total copies of the gene target, typically measured by qPCR (Figure 4B).

**Figure 4.**
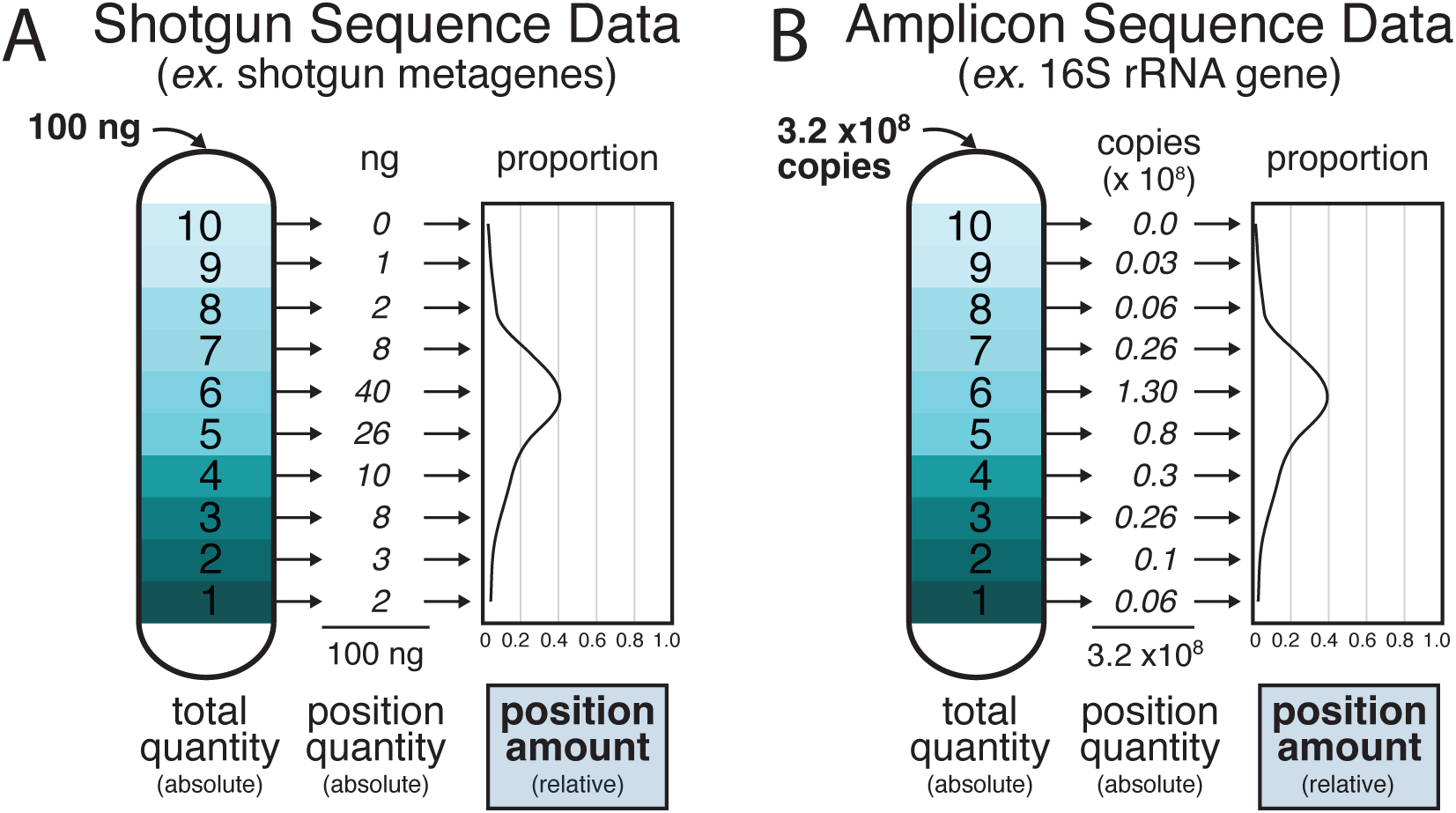
Data on the relative amount of nucleic acid at each gradient position (*gradient_pos_rel_amt*) can be used to calculate taxon-, genome-, or gene-specific AFE using the qSIP method. This value is given as the proportion of nucleic acids at each sequenced gradient position relative to the total nucleic acids available for sequencing. The total amount of nucleic acids depends on the sequencing approach. For shotgun approaches (in A), the *gradient_pos_rel_amt* will correspond to the total mass of nucleic acids (DNA or RNA). For amplicon approaches (in B), the *gradient_pos_rel_amt* will correspond to the total copy number of the targeted gene established via qPCR.

#### SIP methodology

⍰Density gradient formation and the composition of nucleic acids recovered depend on a range of methodological considerations, including rotor type, run speed, run length, gradient medium, fraction volume, and pooling strategy. These methodological details do not fit cleanly into a data standard. However, this information is vital for data interpretation and to explain potential differences among studies in a meta-analysis. The ***sip_method*** field specifies a DOI corresponding to a protocol, article, or data accession in which the complete methodological details have been provided, including any modifications to standard approaches. This type of information can be stored on Protocols.io with a stable DOI [23] when an alternate DOI is not available at the time of archival.

#### Source material identity

SIP sequence data is often complicated by the generation of multiple sequencing products from a single nucleic acid extract (‘sample’). We recommend that depositors specify a unique identifier that links all post-fractionation sequence data to the original (unfractionated) nucleic acid extract. *source_mat_ID* is an existing field in the MIxS framework used to indicate any source from which nucleic acids were sequenced [13]. In MISIP, the ***source_mat_ID*** field is specified as a character string that denotes the unique sample identity for the unfractionated nucleic acid source from which all downstream sequence data originates. We urge the use of Globally Unique IDentifiers (GUID) to maintain the link between the origin of a sample and all downstream measurements. For this, we recommended that a resolvable GUID be used for source samples, which may be the experimental sample (example, soil) or the unfractionated nucleic acid source. Options for resolvable GUIDs include an International Generic Sample Number (IGSN) [24], a BioSample accession number, or an Archival Resource Key (ARK), among others. Notably, if the unfractionated source material was also sequenced, this sample can be included by denoting the *gradient_position* as –1.

#### Isotopolog atom percent

The atom percent of the isotopolog can affect the kinetics of the isotopic labeling of nucleic acids. Low atom percent enrichment of an isotopolog tends to yield more marginal enrichment of nucleic acids and smaller shifts in buoyant density, which can impact sequence data analysis. The ***isotopolog_atom_perc*** field is specified as the atom percent isotope (atom %) of the isotopolog source substrate. If the isotopolog was produced in-house (*ex.* bacterial cellulose from ^13^C-glucose), ensure that the *isotopolog_atom_perc* corresponds with the atom % of the final isotopolog or isotopolog mixture supplied to the system, not the upstream isotopolog used to generate the product.

#### Isotopolog atom position

In cases where an isotopolog is not uniformly labeled, variation in the molecular position of stable isotope atoms can affect the metabolism of isotope into nucleic acids. For example, organisms that preferentially metabolize a functional group, or sidechain, might receive more isotopic label than those that metabolize the whole, or parts, of a partially labeled isotopolog. The ***isotopolog_atom_pos*** field is specified as the International Chemical Identifier (InChI) label, which designates the set of all isotopically enriched atoms present in the isotopolog according to their molecular position [25]. An example of how to generate an InChI label using the InChI open-source chemical structure representation algorithm is provided in the Supplementary Information. The *isotopolog_atom_pos* should only be provided if the isotopolog is a defined compound (*i.e.*, ***isotopolog*** != 0).

#### Isotopolog dose

The concentration of isotopolog (‘dose’) added to the system will also influence the rate and degree of isotopic labeling of nucleic acids. The dose is the mass of isotopolog added per volume of the relevant environmental matrix (*ex.* glucose / total volume soil; CH_4_ / total container volume). The dose should reflect the concentration of isotopolog in the biological system, accounting for the dilution by the environmental matrix, not the concentration of the added isotopolog solution (Figure 5). The dose should also reflect the total cumulative isotopolog added to the system prior to nucleic acid extraction, accounting for multiple additions to the system across time. In cases where the isotopolog is not homogenized within the environmental matrix, the dose should be estimated as the isotopolog concentration in the sample used for nucleic acid extraction. When estimates are too uncertain, or when the concentration of isotopolog in the system is unknown (*ex.* root exudates), no dose should be specified. The ***isotopolog_dose*** field is specified as the final concentration of isotopolog added to the system in parts per million (ppm).

**Figure 5.**
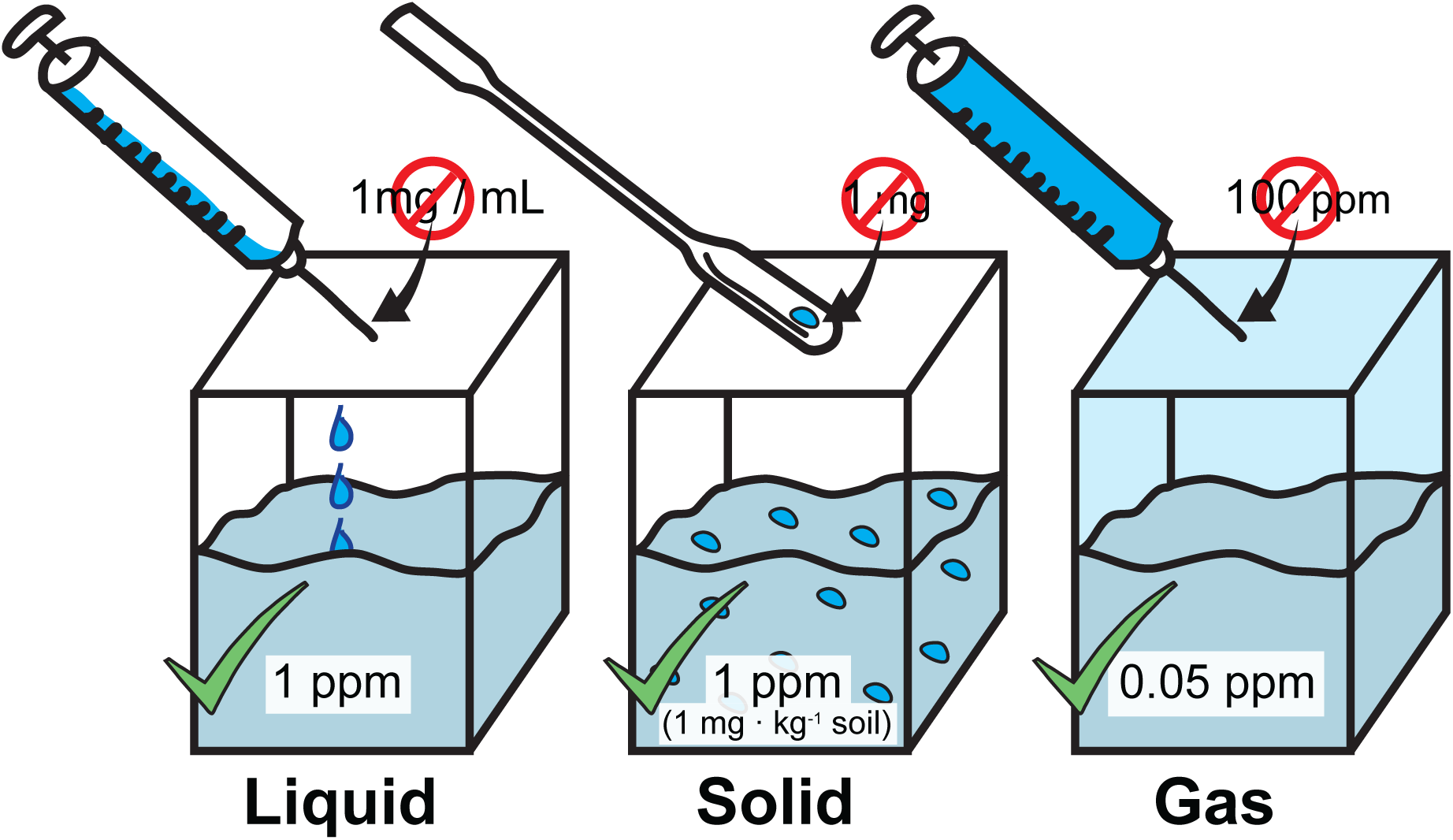
The ***isotopolog_dose*** field provides the concentration, in ppm, of isotopolog in the biological system under study. This field corresponds to the isotopolog exposure of the biological community as measured by the total concentration throughout the system (indicated by the blue fill color), and not the concentration of the amended gas, liquid, or solid material (indicated by the ‘no symbol’). The numbers provided here are intended to serve as examples.

#### Nucleobase atom percent

Bulk measurement of the isotope content of nucleic acids can be used to validate the degree of isotopic enrichment and sequencing-based estimates of AFE. The ***nucleobase_atom_perc*** field is specified as the atom percent (atom %) of isotope in the nucleic acids pool used to generate sequencing data, as previously described [26].

#### Incubation time

The isotopic enrichment of nucleic acids depends on the kinetics of isotopolog metabolism and the fluxes of stable isotope in the experimental system. Over time, isotopic labeling of secondary populations will occur due to access to isotopolog-derived metabolites and biomass. The cross-feeding of stable isotope will produce differences in SIP sequence composition among experiments using the same isotopolog. Thus, providing the incubation time is recommended to aid data interpretation. The ***isotopolog_incu_time*** field is specified as the time in hours (h) from the addition of isotopolog to the end of the incubation period.

#### Internal standard

Internal nucleic acid standards can be used to evaluate gradient formation and normalize gradient position across samples based on the expected range of buoyant densities determined by atom % isotope, GC content, and fragment size [5]. At present, there is no standard methodology for generating or implementing SIP internal standards. Thus, the ***internal_standard*** field specifies a DOI corresponding to a protocol that provides the methodological details, including the sequence composition, isotopic enrichment, and expected buoyant density of any internal standard added to sequencing libraries. This information can be provided on Protocols.io and accessioned with a stable DOI [23]. The internal standard DOI may be the same as the SIP methodology DOI, so long as the SIP methodology DOI fully describes the use of internal standards.

### Accommodating the complexity of SIP experiments

The MISIP data standard specifies the essential information needed to reliably discern the influence of isotopic labeling on the nucleic acid composition of SIP sequence data and improves data reuse by creating a standard vocabulary (Figure 6). These advancements, along with recommendations to guide the collection of other valuable metadata, are sufficient for the majority of SIP experiments. However, the diversity of SIP experimental configurations exceeds the capacity of the MISIP standard to account for every attribute relevant for interpreting a given SIP sequence dataset. Several common, but poorly constrained, experimental attributes were not included in MISIP. For example, the standard does not account for the frequency and timing of multiple doses of isotopolog in a pulse-chase type of experiment, the homogeneity of isotopolog during the incubation, or whether an experimental system is open or closed to the environment, which can alter isotopic labeling due to the influx of unlabeled isotopologs or efflux of stable isotope label. The standard also does not capture kinetics data relevant to the metabolic activity governing the isotopic enrichment of nucleic acids, such as respiration (^13^CO_2_) or other metabolic byproducts [27], that provide information about the latency of isotope assimilation and cross-feeding. To address these limitations, and to optimally curate SIP metadata for reuse, we advise depositors to use the MISIP standard to guide data collection in planning and performing a SIP experiment. Once data has been collected, we advise depositors to complete the MISIP-MIMS and/or MISIP-MIMARKS checklists in tandem with writing a description of their methods [18]. This will help ensure the relevant metadata not captured by the checklists will be included in methodological descriptions available elsewhere in the public record.

**Figure 6.**
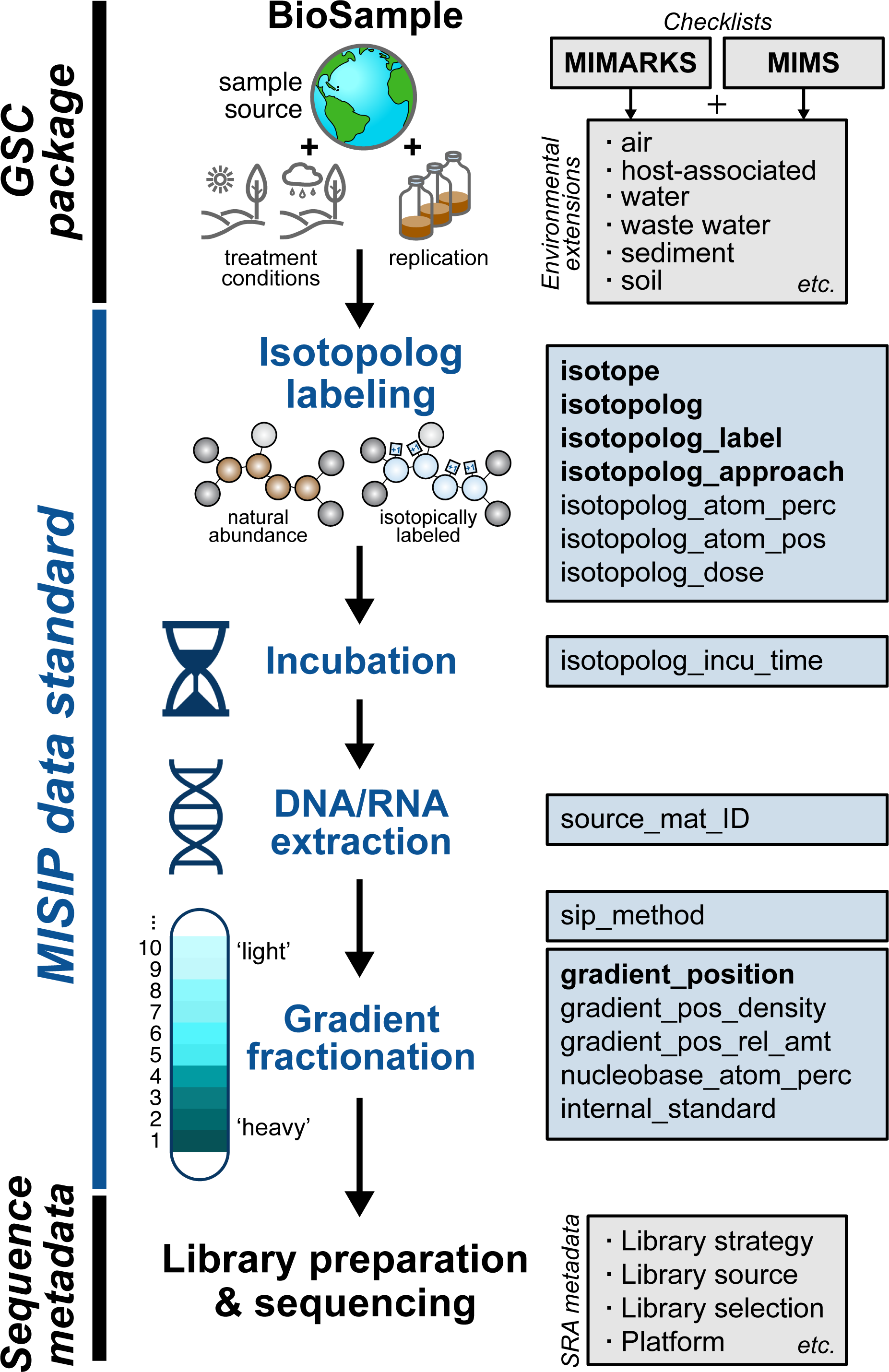
The MISIP standard captures metadata at several stages during a SIP experiment. This schematic provides an overview of the typical workflow of DNA/RNA-SIP experiment and the SIP-specific metadata (highlighted in blue boxes) that can be collected at each step. The related upstream and downstream metadata required for sequence data archival covered by other data standards is highlighted in black boxes.

The fields described in the MISIP standard are designed to be combined with at least one other MIxS checklist (MIMS or MIMARKS) plus an environmental extension. The MISIP data standard, alone, does not retain the full breadth of metadata necessary for interpretation and reuse without additional methodological information, such as nucleic acid extraction methods (*nucl_acid_ext*) or the primers used to generate gene marker data (*pcr_primers*), and other contextual information in an environmental extension, such as the location sampled (*lat_lon*), chemical descriptions of the source (*ex. carb_nitro_ratio*), and other chemicals administered (*chem_administration*). We encourage researchers to use the existing fields in MIMS, MIMARKS, and MIxS environmental extensions to describe environmental source and sequence preparation more completely.

### Advice on the reuse of SIP sequence data

The MISIP standard is designed to encourage the reuse of SIP sequence data. To that end, we offer brief advice to assist in reanalyzing existing datasets. The primary focus of the analysis should be to contrast sequence data from equivalent gradient positions between paired samples supplied with either isotopically labeled or natural abundance isotopolog. We advise <u>against</u> making comparisons within a density gradient using isotopically labeled sequence data without the context of natural abundance data. Compositional differences may be driven by GC content and other factors, which are best controlled by comparison to sequence data from the paired natural abundance sample. That said, the inclusion of paired natural abundance samples is not required by MISIP and, in cases where it is missing, it may be possible to model the gradient distribution of natural abundance nucleic acid fragments [21, 28].

Although the MISIP standard aims to reduce error from mismatched gradient fractions, there is always a possibility of human error. For this reason, the inclusion of buoyant density information is highly recommended. However, in cases where density information is not provided, we advise a user to iterate their analysis using a ‘sliding window approach,’ where adjacent gradient positions are combined to offset small variations in density at any one position.

## Conclusions

Over two decades, the global SIP community has worked together to establish fundamental methods that enable researchers to connect biogeochemical and microbial processes and leverage the omics analytical toolkit. The current movement toward better standardization of ‘omics’ metadata promises to expand the linkages between diverse data types relevant to SIP sequence data. A common stable isotope source offers an inherent signal from which to integrate diverse omics data types (*i.e.*, proteomic, metabolomic, etc.). Such an approach will provide a systems-level view of biological processes but will also require flexible frameworks to ingest different SIP datatypes and metadata. The MISIP standard was developed to provide a foundation for these efforts by formalizing the minimum metadata requirements for any SIP-derived sequence and to formalize a common vocabulary for SIP metadata. By providing a shared vocabulary to guide metadata entry and validation, MISIP can assist in the development of bioinformatic software for SIP data analysis and the design of SIP data intake on platforms that extend a FAIR framework [12] to downstream data analysis, ensuring SIP continues to deepen our understanding of the activity of biological communities.

## Acknowledgements

We would like to thank Kaitlyn Johnson and Payton Taylor for their help curating the list of SIP studies. We would like to thank Dr. Samuel Barnett for providing valuable feedback on the standard and the manuscript and Dr. Rex Malmstrom for feedback on the proposed standard. We are extremely grateful for the guidance received from members of the GSC, specifically Mark Miller, MSc, Dr. Lynn Schriml, and Dr. Chris Hunter. Partial funding for RCW was provided by the USDA National Institute of Food and Agricultural Hatch grant (IND90024429).

