## Supplemental File 1 for "A data standard for the reuse and reproducibility of any stable isotope probing-derived nucleic acid sequence (MISIP)"

**Supplementary Information**

This information corresponds to the article by Simpson *et al.*, 2023 entitled “A data standard for the reuse and reproducibility of any stable isotope probing-derived nucleic acid sequence (MISIP).”

*Generating International Chemical Identifier (InChI) Label*

The InChI label is a machine-readable format that specifies the exact position of all atoms in a molecule, including the isotope number (at the end of the label). The label is a stable identifier supported by the IUPAC. At the time of publication, the following freely available software tools could be used to generate an InChI label using visualizations to guide the appropriate designation of isotopic labeling. The same tools can be used to visualize an InChI label as a molecular structure. We will demonstrate how to obtain an InChI for the commercially available ^13^C-labeled L-DOPA (IUPAC: (S)-2-Amino-3-(3,4-dihydroxyphenyl)propanoic acid).

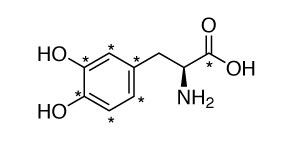

**Step 1**: Install the Marvin Sketch software developed by Chemaxon <https://chemaxon.com/marvin>. The software is available for non-commercial use under an academic license.

**Step 2**: In Marvin Sketch, select ‘Import Name’ and specify the IUPAC name for your isotopolog compound.

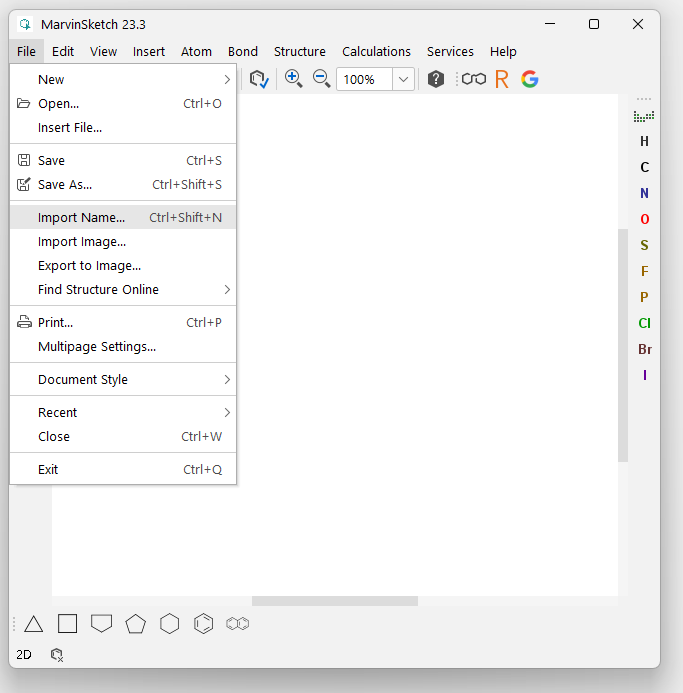

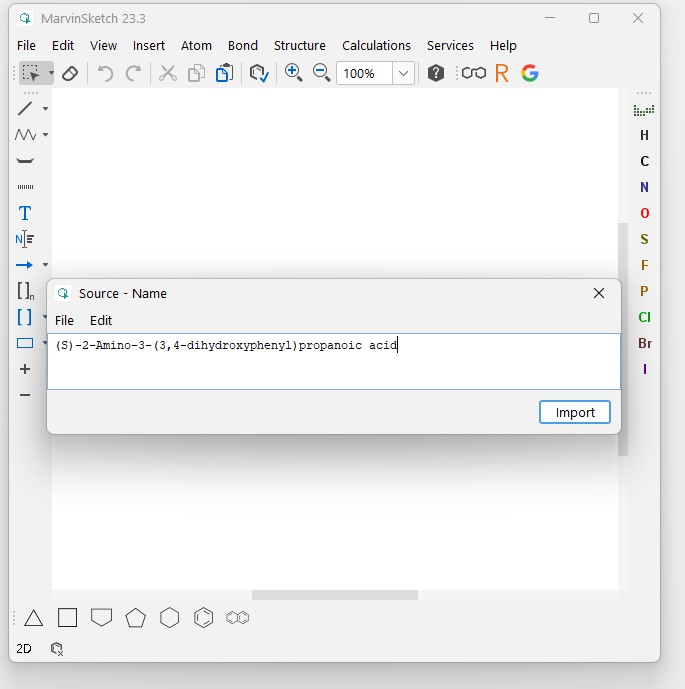

**Step 3**: The molecular structure will be displayed. Select each atom that is isotopically labeled using your cursor. Once selected, click ‘Atom,’ then ‘Isotope,’ and then the correct isotope.

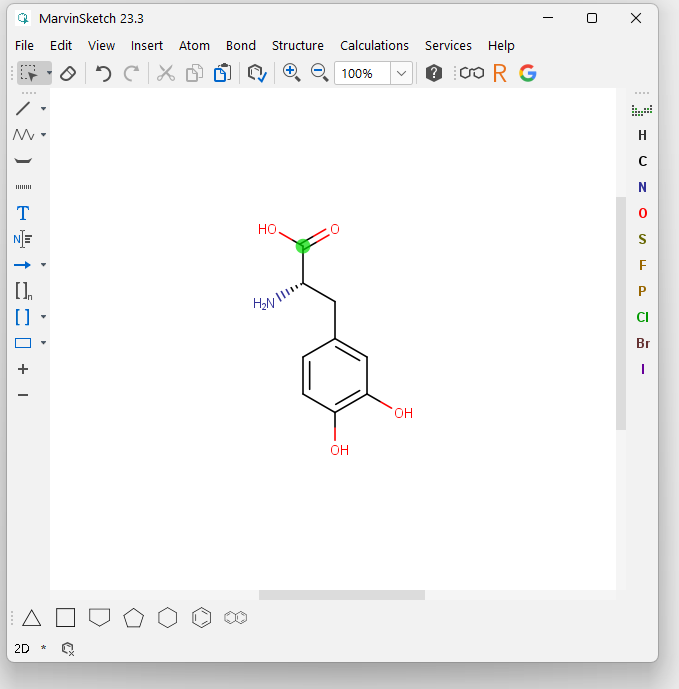

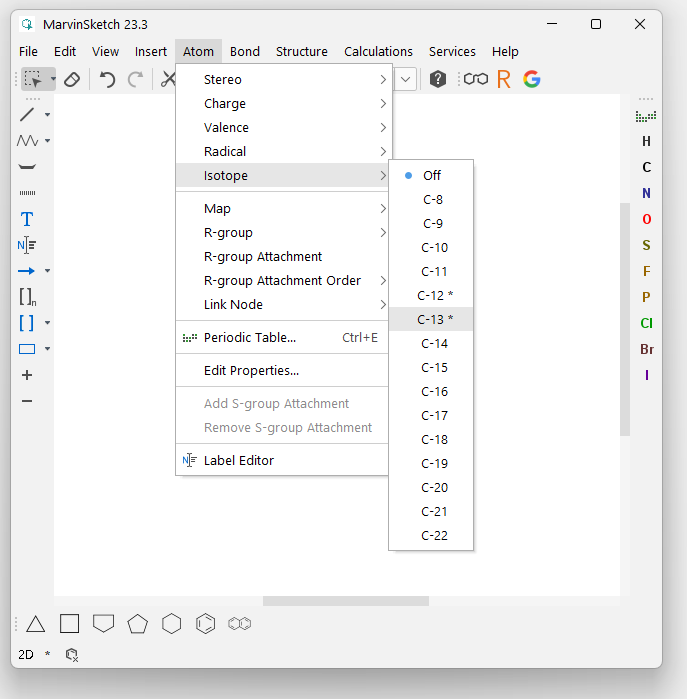

**Step 4**: Repeat until all atoms are labeled, then select ‘File’ and ‘Save As’ from the menu bar. Save the file as either an MDL SDfile (.sdf) or Tripos SYBYL Molfile (.mol).

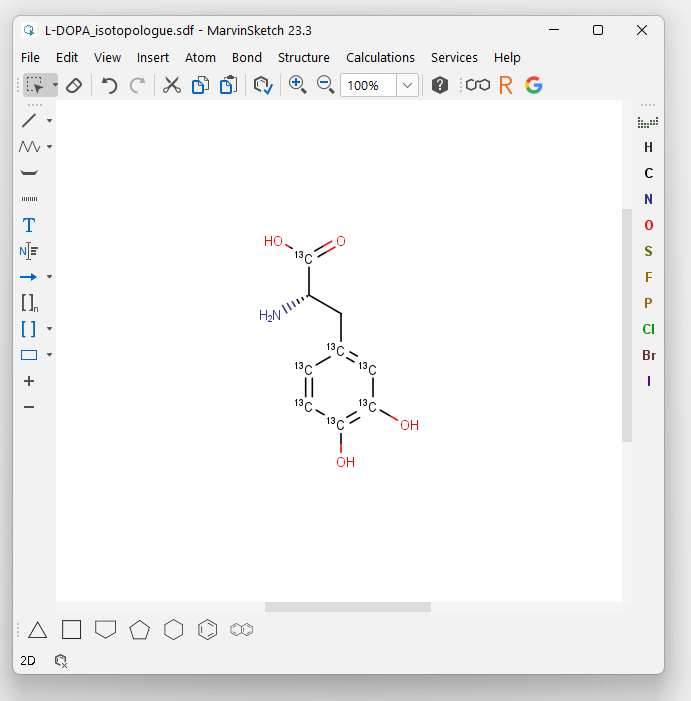

**Step 5**: Either (1) use the software ‘OpenBabel’, available online at http://www.cheminfo.org/Chemistry/Cheminformatics/FormatConverter/index.html to convert the .sdf or .mol to the InChI format, or (2) download the InChI software binaries from the InChI Trust (available for Linux or Windows OS). Start the software using the executable file. Once running, select ‘Open’ and navigate to the file outputted from Marvin Sketch. The InChI label for your isotopolog will be displayed in the dialogue box beneath the molecular structure. Double check that all isotopically labeled atoms are correctly assigned in your structure, then copy and paste the InChI into your MISIP spreadsheet. In our example this value would be: “1S/C9H11NO4/c10-6(9(13)14)3-5-1-2-7(11)8(12)4-5/h1-2,4,6,11-12H,3,10H2,(H,13,14)/t6-/m0/s1/i1+1,2+1,4+1,5+1,7+1,8+1,9+1”

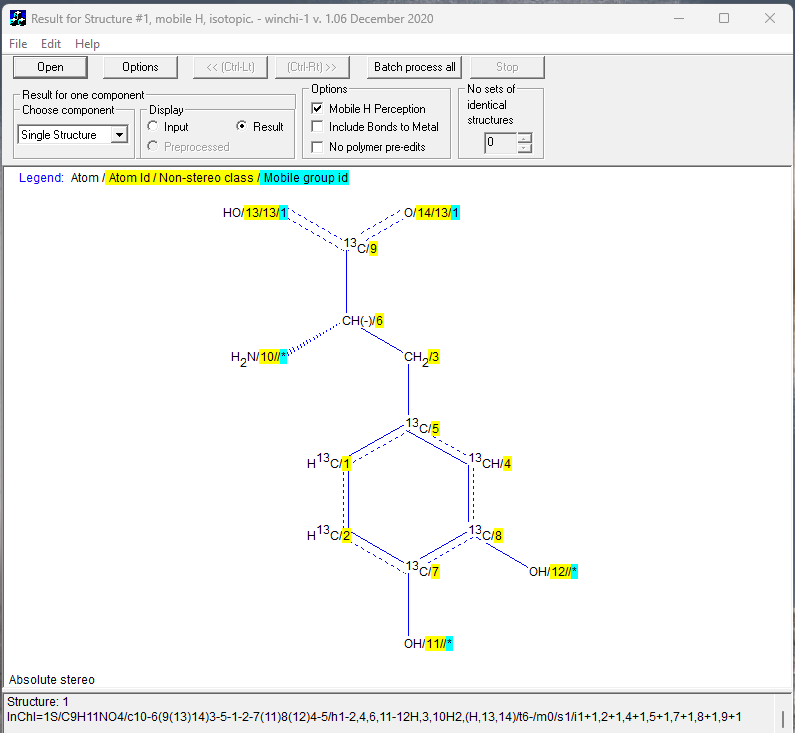

For your convenience, listed below are some isotopologs commonly used in DNA/RNA-SIP and their InChI label:

| ^18^O water | 1S/H2O/h1H2/i1+2 |
| --- | --- |
| ^13^C carbon dioxide | 1S/CO2/c2-1-3/i1+1 |
| ^13^C bicarbonate | 1S/CH2O3/c2-1(3)4/h(H2,2,3,4)/p-1/i1+1 |
| ^13^C methane | 1S/CH4/h1H4/i1+1 |
| ^13^C methanol | 1S/CH4O/c1-2/h2H,1H3/i1+1 |
| Uniformly ^13^C acetate | 1S/C2H4O2/c1-2(3)4/h1H3,(H,3,4)/p-1/i1+1,2+1 |
| Uniformly ^13^C glucose | 1S/C6H12O6/c7-1-3(9)5(11)6(12)4(10)2-8/h1,3-6,8-12H,2H2/t3-,4+,5+,6+/m0/s1/i1+1,2+1,3+1,4+1,5+1,6+1 |
| Ring-labeled ^13^C vanillin | 1S/C8H8O3/c1-11-8-4-6(5-9)2-3-7(8)10/h2-5,10H,1H3/i2+1,3+1,4+1,6+1,7+1,8+1 |
| Ring-labeled ^13^C toluene | 1S/C7H8/c1-7-5-3-2-4-6-7/h2-6H,1H3/i2+1,3+1,4+1,5+1,6+1,7+1 |
| Uniformly ^13^C toluene | 1S/C7H8/c1-7-5-3-2-4-6-7/h2-6H,1H3/i1+1,2+1,3+1,4+1,5+1,6+1,7+1 |
| ^13^C urea | 1S/CH4N2O/c2-1(3)4/h(H4,2,3,4)/i1+1 |
| Uniformly ^15^N urea | 1S/CH4N2O/c2-1(3)4/h(H4,2,3,4)/i2+1,3+1 |
| Uniformly ^15^N dinitrogen | 1S/N2/c1-2/i1+1,2+1 |
