## Supplementary material for "A data standard for the reuse and reproducibility of any stable isotope probing-derived nucleic acid sequence (MISIP)": Figure S1

**Figure S1**. An overview of the quality of SIP metadata archived at the SRA from various environments. SIP studies accessioned in PubMed with a Sequence Read Archive containing more than five samples were evaluated for their metadata quality. SRAs which reported the isotopolog, isotopolog label status, and gradient position met the "Minimum" (Min) requirements, while those that reported at least one of these items were "Insufficient", and those that reported none were categorized as "None."

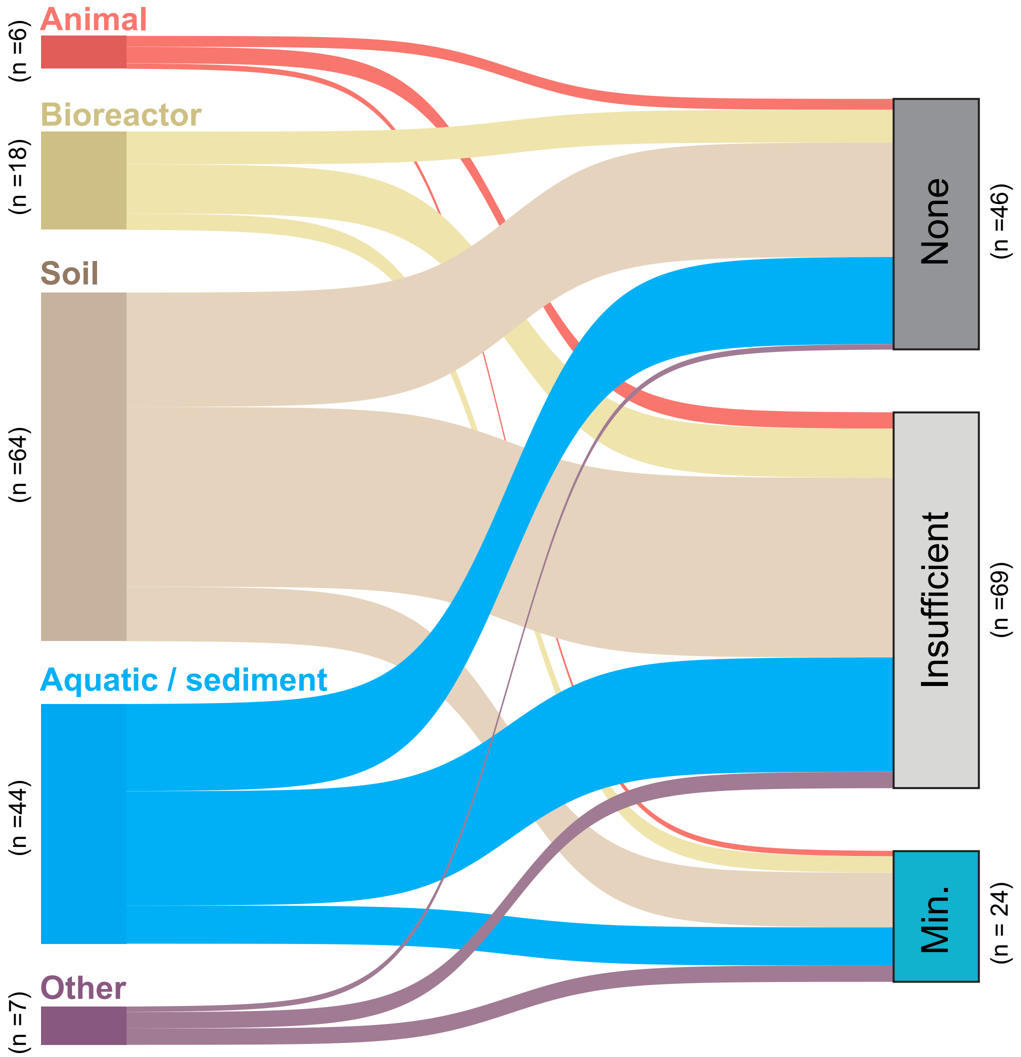
